# The anaerobic gut fungal community in ostriches (*Struthio camelus*)

**DOI:** 10.1101/2025.03.28.646006

**Authors:** Julia Vinzelj, Kathryn Nash, Adrienne L. Jones, R. Ty Young, Casey H. Meili, Carrie J. Pratt, Yan Wang, Mostafa S. Elshahed, Noha H. Youssef

**Affiliations:** Department of Microbiology and Molecular Genetics, Oklahoma State University, Stillwater, OK, U.S.A.; Department of Biological Sciences, University of Toronto Scarborough, Toronto, ON, Canada

**Author notes:** Corresponding author. Address: 1110 S. Innovation Way, Stillwater, OK, 74074.

## Abstract

Anaerobic gut fungi (AGF; *Neocallimastigomycota*) are essential for plant biomass degradation in herbivores. While extensively studied in mammals, information regarding their occurrence, diversity, and community structure in non-mammalian hosts remains sparse. Here, we report on the AGF community in ostriches (*Struthio camelus*), herbivorous, flightless, hindgut fermenting members of the class *Aves* (birds). Culture-independent diversity surveys of fecal samples targeting the D2 region of the large ribosomal subunit (28S rRNA) revealed a uniform community with low alpha diversity. The community was mostly comprised of sequences potentially representing two novel species in the genus *Piromyces,* and a novel genus in the *Neocallimastigomycota*. Sequences affiliated with these novel taxa were absent or extremely rare in datasets derived from mammalian and tortoise samples, indicating a strong pattern of AGF-host association. One *Piromyces* strain (strain Ost1) was successfully isolated. Transcriptomics-enabled molecular dating analysis suggested a divergence time of ≈ 30 Mya, a time frame in line with current estimates for ostrich evolution. Comparative gene content analysis between strain Ost1 and other *Piromyces* species from mammalian sources revealed a high degree of similarity. Our findings expand the range of AGF animal hosts to include members of the birds (class *Aves*), highlight a unique AGF community adapted to the ostrich alimentary tract, and demonstrate that – like mammals – coevolutionary phylosymbiosis (i.e. concurrent evolution of AGF and their animal hosts) plays a central role in explaining current AGF distribution patterns in *Aves*.

## Introduction

Anaerobic gut fungi (AGF) are a clade of basal, zoospore producing fungi belonging to the phylum *Neocallimastigomycota* within the subkingdom *Chytridiomyceta*^1^. They inhabit the digestive tracts of herbivores, crucially aiding in the degradation of plant material and its fermentation.^2,3^ To date, 22 genera of AGF have been described^4,5^, though large-scale culture-independent surveys predict at least twice as many still uncultured^6^. AGF were originally isolated from placental mammals^7^, and their occurrence and diversity have been extensively studied in domesticated mammalian hosts (e.g., cows, goats, sheep, and horses) owing to their economic importance and ease of sampling. Large-scale culture-independent studies, however, have also identified AGF in many additional wild mammalian^6,8^, marsupial^9^, and non-mammalian hosts such as green iguanas^10^, and tortoises^11^. The recent isolation of novel strains belonging to basal clades of AGF from tortoises highlights the underexplored scope of diversity and host range of AGF^5^.

It is currently unclear what exactly defines the AGF ecological niche, though factors such as host phylogeny, herbivory, prolonged feed retention time, and the presence of dedicated fermentation sites within the digestive tract have been proposed as key determinants. Among these, host phylogeny has been shown to have a greater impact on AGF community composition than diet or other environmental factors^6,10,12^. AGF are slow-growers and tend to adhere to plant material^13^, suggesting that their survival in a competitive environment might, in part, be dependent on longer retention times. This aligns with the mean feed retention times found in ruminants (43-75 hours)^14^ and mammalian hindgut fermenters (24-47 hours)^14–16^, as compared to the retention time found in carnivores.^17^ Tortoises exhibit the longest retention time (7-14 days)^18–20^ of all AGF hosts identified so far. To the best of our knowledge, no comprehensive study has defined the ecological niche of AGF in more detail, though reports exist on the absence of AGF in omnivores like pigs^21^ and dogs.^22^

Birds (class *Aves*), a lineage of warm-blooded, non-mammalian vertebrates within the clade *Sauropsida*, exhibit great ecological and physiological diversity. While most extant bird species are omnivorous, an estimated 2% thrive on mostly herbivorous diets.^23,24^ Many of those herbivorous birds compensate for the low digestibility of plant matter by increasing food intake and shortening gastrointestinal retention times^23–25^, adaptations that are potentially unfavorable for the establishment of AGF. Ostriches (genus *Struthio*), however, represent an exception within *Aves*. As large, herbivorous, flightless members of the *Palaeognathae* (an infraclass that also includes rheas, cassowaries, emus, and kiwis), they primarily consume grasses, shrubs, and succulents.^25,26^ They are known to be apt lignocellulose degraders, degrading up to 60% of grasses and leaves eaten.^27^ They are equipped with highly specialized gastrointestinal adaptations, including the gizzard filled with grit for mechanical disruption and storage of plant biomass, and large, compartmentalized sacculated ceca with an elongated and partly sacculated colon as main fermentation sites.^28^ The retention time in ostriches (30-40 hours^29,30^) resembles that of mammalian hindgut fermenters. Additionally, the ceca and colon are highly efficient in the absorption of water from the digesta, probably an adaptation to the arid environment in which ostriches evolved.^29^

We hypothesized that the alimentary tract architecture and nutritional preferences of ostriches would allow for the establishment and propagation of AGF as a component of their gut microbiome. To test this hypothesis, we examined the occurrence, diversity, and community structure of AGF communities in ostrich fecal samples using a combination of culture-independent and culture-based surveys. Our results highlight the novelty of AGF taxa encountered in ostriches as well as the differences and similarities compared to their mammalian counterparts. The ecological and evolutionary implications of these findings are discussed.

## Materials and Methods

### Samples

Ostrich fecal samples (n = 13) were collected between 2020 and 2022 from the Oklahoma City Zoo, as well as private ranches in Oklahoma and Texas, U.S.A. (Table S1). All samples were obtained from adult ostriches, mostly fed a pellet diet composed of corn, soy, and pre-mixed commercially sold ostrich feed. The samples were obtained shortly after defecation and collected in 15- or 50-ml Falcon tubes that were placed on ice during transfer to the laboratory, where they were stored at −20°C.

### DNA extraction and amplification

DNA extraction from fecal samples was performed using the DNeasy Plant Pro kit (Qiagen®, Germantown, Maryland, U.S.A.) according to the manufacturer’s instructions. For detection and characterization of the AGF community, the primer pair AGF-LSU-EnvS For and AGF-LSU-EnvS Rev with Illumina overhang adaptors was used^6,22^. DreamTaq 2X master mix (Life Technologies, Carlsbad, California, U.S.A.) was used for amplification of the D2 region of the LSU rRNA according to the manufacturer’s instructions, and the PCR protocol for all reactions (excluding indexing) was the same as previously described.^9^ Each PCR run also included a non-template control to monitor potential contamination. Given that fecal samples are known to produce DNA extracts containing PCR inhibitors^31^, additional efforts to obtain amplicons from samples initially showing negative PCR amplification included varying the DNA concentrations and DNA to primer ratio.

### Sequencing and sequence processing

PCR cleaning, indexing, and pooling were conducted as previously described.^9^ Pooled libraries were sequenced either at the University of Oklahoma Clinical Genomics Facility (Oklahoma City, Oklahoma, U.S.A.) using the MiSeq platform and the 300 bp PE reagent kit, or at the Oklahoma State University One Health Innovation Foundation (Stillwater, Oklahoma, U.S.A.) using the NextSeq platform and the 300 bp PE reagent kit. Sequence quality control was conducted as previously described.^6^ Coverage values were calculated using the command *phyloseq_coverage* in the R package metagMisc.^32,33^

A two-tier approach, as detailed before^6^, was used to assign sequences to previously described genera and candidate genera and to identify novel AGF genera. Genus-level assignments were used to build a shared file (using the mothur commands *phylotype* and *make.shared*), which was then utilized as an input for downstream analysis. Confirmatory amplification and sequencing of the longer D1-D2 LSU fragment (∼ 700 bp) was conducted on select samples using the primers NL1F (5’-GCATATCAATAAGCGGAGGAAAAG-3’) and GG-NL4 (5’-TCAACATCCTAAGCGTAGGTA-3’) primers, and PacBio sequencing as previously described.^6^

Sequences identified as members of the genus *Piromyces* were further binned into species-level operational taxonomic units (OTUs) by assessing percentage divergence patterns to reference cultured *Piromyces* sequences. A cutoff of 3% divergence was used, since it reflects the average divergence between various currently described *Piromyces* species.^34,35^ For ecological distribution analysis, representative sequences of the three most encountered species-level OTUs in ostrich fecal samples were used to query their occurrence in prior broad host diversity surveys.^6,9,11,22^

### Alpha diversity

The R package phyloseq (v 1.50.0)^32^ was used to calculate alpha diversity estimates (Observed, Shannon, Simpson, and Inverse Simpson diversity indices) using the command *estimate_richness*. Alpha diversity estimates from ostrich datasets were compared to estimates from a subset of mammalian counterparts (25 cattle, 25 goats, 25 sheep, 24 deer, 25 horses) included in a recent study of the mammalian AGF mycobiome^6^, as well as to datasets from tortoises obtained in another study (n = 11 tortoises).^11^ The two-sided Wilcoxon signed rank test for pairwise comparison of means was used to examine the effect of animal species, family, and class on alpha diversity estimates.

### Community structure

The phylogenetic similarity-based weighted Unifrac index (calculated using the *ordinate* command in the phyloseq R package) was used to construct principal coordinate analysis (PCoA) ordination plots using the function *plot_ordination* in the phyloseq R package. The AGF community structure in ostriches was compared to that found in mammalian and reptilian hosts using the same data set used for alpha-diversity comparisons (see above). To partition the dissimilarity among host factors (animal species, family, class, and gut type), we used PERMANOVA tests (using the command *adonis* in the R package vegan, v2.6-8).^33^ Factors that significantly affect the AGF community structure were identified using the F-statistics and p-values, and the percentage variance explained by each factor was calculated as the percentage of the sum of squares of each factor to the total sum of squares.

To identify AGF genera differentially abundant in ostriches, the genus-level shared file created in mothur was used to calculate both linear discriminant analysis (LDA) effect size (LEfSe) and Metastats. Genera with calculated LDA scores and/or significant Metastats p-values were considered differentially abundant. Further, to identify AGF community members responsible for the observed community structure in ostriches versus mammalian or reptilian species, Bray-Curtis index values (calculated using the *ordinate* command in the phyloseq R package) were used to construct double principal coordinate analysis (DPCoA) ordination plots using the function *plot_ordination* in the phyloseq R package. To assess ostrich–AGF genera associations, we calculated global phylogenetic signal statistics (Abouheif’s Cmean, Moran’s I, and Pagel’s Lambda) using the *phyloSignal* command in the phylosignal R package (v. 1.3.1)^36^, as well as the Local Indicator of Phylogenetic Association (LIPA) using *lipaMoran* command in the phylosignal R package.

### Phylogenetic tree construction

The phylogenetic position of novel AGF genera and species was evaluated by constructing maximum likelihood phylogenetic trees in FastTree^37^ based on the MAFFT-generated multiple sequence alignment of the D2 LSU rRNA sequences of the novel taxa to the D1-D2 region of all previously reported cultured and uncultured AGF genera as references (version 2.0, https://anaerobicfungi.org/databases).

### Enrichment and isolation attempts for AGF from ostrich feces

Enrichments were set up in an anaerobic chamber (Coy Laboratories, Grass Lake, Michigan, U.S.A.) as previously described^7^ using different substrates (Table 1) in either rumen fluid cellobiose (RFC)^38^ or rumen fluid free medium (RFF).^39^ To obtain pure cultures, multiple rounds of sub-cultivation and roll tubes were conducted as previously described.^5^ Identity of the isolates was determined by amplifying and Sanger-sequencing the D1-D2 region of the LSU rRNA gene using primers NL1F and NL4R (5’-GGTCCGTGTTTCAAGACGG-3’). Sequencing was conducted at the Oklahoma State University Biochemistry and Molecular Biology Core Facility.

**Table 1.** Enrichments/isolation attempts with ostrich fecal samples targeting *Neocallimastigomycota*.

| Species name | Number of enrichments set up | Number of successful enrichments | Media used | Incubation temperatures (°C) | Samples used |
| --- | --- | --- | --- | --- | --- |
| *Piromyces* sp. Ost1 | 7 | 7 | RFC+SG, RFF + SG | 39, 41, 43 | OS_F1, OS_F2 |
| *Piromyces* sp. Ost2 | 10 | 0 | RFC+SG, RFF + SG | 39, 41 | TY06, TY07, TY08 |
| JV1 | 6 | 2 | RFC + C | 39, 41 | OS_F2 |

<sup>1</sup>RFC = rumen-fluid cellobiose medium,
<sup>2</sup>RFF = rumen fluid free medium,
<sup>3</sup>+ SG = adding switchgrass as C-source,
<sup>4</sup>+ C = adding cellulose as C-source

### Transcriptomic sequencing and timing

*Piromyces* sp. Ost1 cultures were grown in RFC medium to late exponential phase, early stationary phase (five days), vacuum filtered, and total RNA was extracted using the Macherey-Nagel™ NucleoSpin™ RNA Mini kit according to the manufacturer’s instructions. *Piromyces* sp. Ost1 RNA-seq was conducted on an Illumina NextSeq 2000 platform using a 2×150 bp paired-end library at the One Health Innovation Foundation lab at Oklahoma State University. RNA-Seq reads were quality trimmed and *de novo* assembled using Trinity (version 2.6.6)^40^ with default parameters. Transcripts were clustered with an identity parameter of 95 % (–c 0.95) using CD-HIT^41^ to remove redundancy.

Remaining transcripts were then used for peptide and coding sequence predictions using TransDecoder (version 5.0.2) (Haas, B.J. https://github.com/TransDecoder/TransDecoder) with a minimum peptide length of 100 amino acids. Gene content of *Piromyces* sp. Ost1 transcriptome was compared to nine previously sequenced *Piromyces* transcriptomes^35,42–44^, all isolated from mammals and belonging to six different putative *Piromyces* species. Comparative gene content analysis was carried out via classification of all predicted peptides from all transcriptomes against COG (via BLASTp comparisons against the most updated database at https://ftp.ncbi.nih.gov/pub/COG/COG2020/data/), KOG (via blastp comparisons against the most updated database at https://ftp.ncbi.nih.gov/pub/COG/KOG/), and KEGG classification (by running GhostKOALA^45^ search on the predicted peptides) schemes.

To examine the CAZymes production potential of *Piromyces* sp. Ost1 compared to other mammalian isolates (belonging to other *Piromyces* species, as well as to other AGF genera, n=53)^6,42–44,46,47^, as well as tortoise isolates (*Testudinimyces* and *Astrotestudinimyces*, n =7)^11^, we predicted the overall CAZyme content (using run_dbcan4 (https://github.com/linnabrown/run_dbcan) to identify glycoside hydrolases (GHs), polysaccharide lyases (PLs), carbohydrate esterases (CEs), alpha amylases (AAs), and carbohydrate-binding motifs (CBMs).

### Phylogenomic analysis and molecular dating

We used the predicted peptides from the *Piromyces* sp. Ost1 transcriptome, as well as the 60 available AGF transcriptomes^6,11,35,42–44,46,47^ for phylogenomic analysis and molecular timing of evolutionary divergence, as previously described.^6,11,35^ Five *Chytridiomycota* genomes (*Chytriomyces* sp. strain MP 71, *Entophlyctis helioformis* JEL805, *Gaertneriomyces semiglobifer* Barr 43, *Gonapodya prolifera* JEL478, and *Rhizoclosmatium globosum* JEL800) were used as outgroups and to provide calibration points.

We used the ‘fungi_odb10’ dataset, including 758 phylogenomic markers for kingdom *Fungi*^48^, for our analysis. Profile hidden Markov models (HMMs) of these markers were previously created and used for previous AGF phylogenomic studies.^6,11,35^ HMMs were used to identify homologues in all AGF transcriptomes, as well as the five *Chytridiomycota* genomes using HMMER3 (http://hmmer.org/). Markers identified with conserved homologs in all datasets were aligned and concatenated for subsequent phylogenomic analyses. IQ-TREE^49^ was used to find the best-fit substitution model and to reconstruct the phylogenetic tree with the maximum-likelihood approach. PartitionFinder (v 2.1.1)^50^ was used to group the refined alignment and to assign each partition with an independent substitution model. All partition files, along with their corresponding models, were then imported into BEAUti (v 1.10.4)^51^ for conducting Bayesian and molecular dating analyses. Two calibration priors were set: a direct fossil record of *Chytridiomycota* from the Rhynie Chert (407 Mya) and the emergence time of *Chytridiomycota* (573 to 770 Mya as 95% HPD). We used the Birth-Death incomplete sampling tree model for interspecies relationship analyses. Unlinked strict clock models were used for each partition independently. Three independent runs (30 million generations each) were performed with a default burn-in (10%). Tracer (v1.7.1)^52^ was then used to confirm that a sufficient effective sample size (ESS > 200) was obtained. Finally, TreeAnnotator (v1.10.4)^51^ was used to compile the maximum clade credibility (MCC) tree.

### Sequence and data deposition

Illumina and RNA-seq reads were deposited in NCBI SRA under BioProject accession number PRJNA1231060. Clone sequences of the D1-D2 region of the LSU rRNA from the *Piromyces* sp. Ost1 isolate were deposited in GenBank under accession numbers PV213533-PV213569. PacBio sequence representatives of the *Piromyces* sp. Ost2 and candidate genus JV1 were deposited in GenBank under accession numbers PV226234 and PV226233, respectively.

## Results

### Occurrence and AGF community composition in ostriches

A total of 342,691 high-quality AGF-affiliated D2-LSU sequences (average per sample: 26,361) were obtained (Table S1). High coverage values indicated that most of the genus-level diversity was captured in all samples (Table S1).

Phylogenetic analysis indicated that the AGF community in ostriches displayed a high level of similarity and was dominated by sequences affiliated with two genera. Sequences affiliated with the genus *Piromyces* constituted >92% of the community in 11/13 samples and roughly half (44.6 and 51.5%) of the community in the remaining two samples (Figure 1A, Table S1). The majority of *Piromyces* sequences clustered into two species-level operational taxonomic units (OTUs) (Figure 1B). Both OTUs were phylogenetically distinct from previously named *Piromyces* for which D1-D2 LSU sequence data is currently available (*P. finnis*^53^, *P. rhizinflata*^54^, and *P. communis*^55^), as well as previously reported and yet to be named isolates [*Piromyces* sp. A1^56^, *Piromyces* sp. B4^56^, *Piromyces* sp. NZB19^57^, *Piromyces* sp. PR1 (unpublished, GenBank accession number JN939159), and *Piromyces* sp. Axs (unpublished, GenBank accession number PV351789)]. As such, the OTUs found in ostriches represent two putative novel *Piromyces* species, for which Ost1 and Ost2 designations are proposed (Figure 1C). *Piromyces* sp. Ost1 exhibited 96.55% sequence similarity to its closest relative (*Piromyces* sp. A, GenBank accession MT085679.1), while *Piromyces* sp. Ost2 exhibited 95.25% sequence similarity to its closest relative (*Piromyces communis* Clone P, GenBank accession ON619893.1) (Table 1). Assessment of the occurrence of these species in prior AGF-focused culture-independent diversity surveys^8,11,14,24^ showed that they are either absent or only a minor part of the community in the hosts investigated. For example, *Piromyces* sp. Ost1 was completely absent in all mammalian fecal samples examined in Meili *et al*.^6^, absent in 12 out of 15 samples examined in Young *et al*.^22^, and extremely rare in tortoise (present in one out of 11 samples, constituting 0.02% of total sequences from tortoises)^11^ and marsupial (present in two out of 61 samples, constituting 0.001% of total sequences from marsupials)^9^ samples.

**Figure 1.**
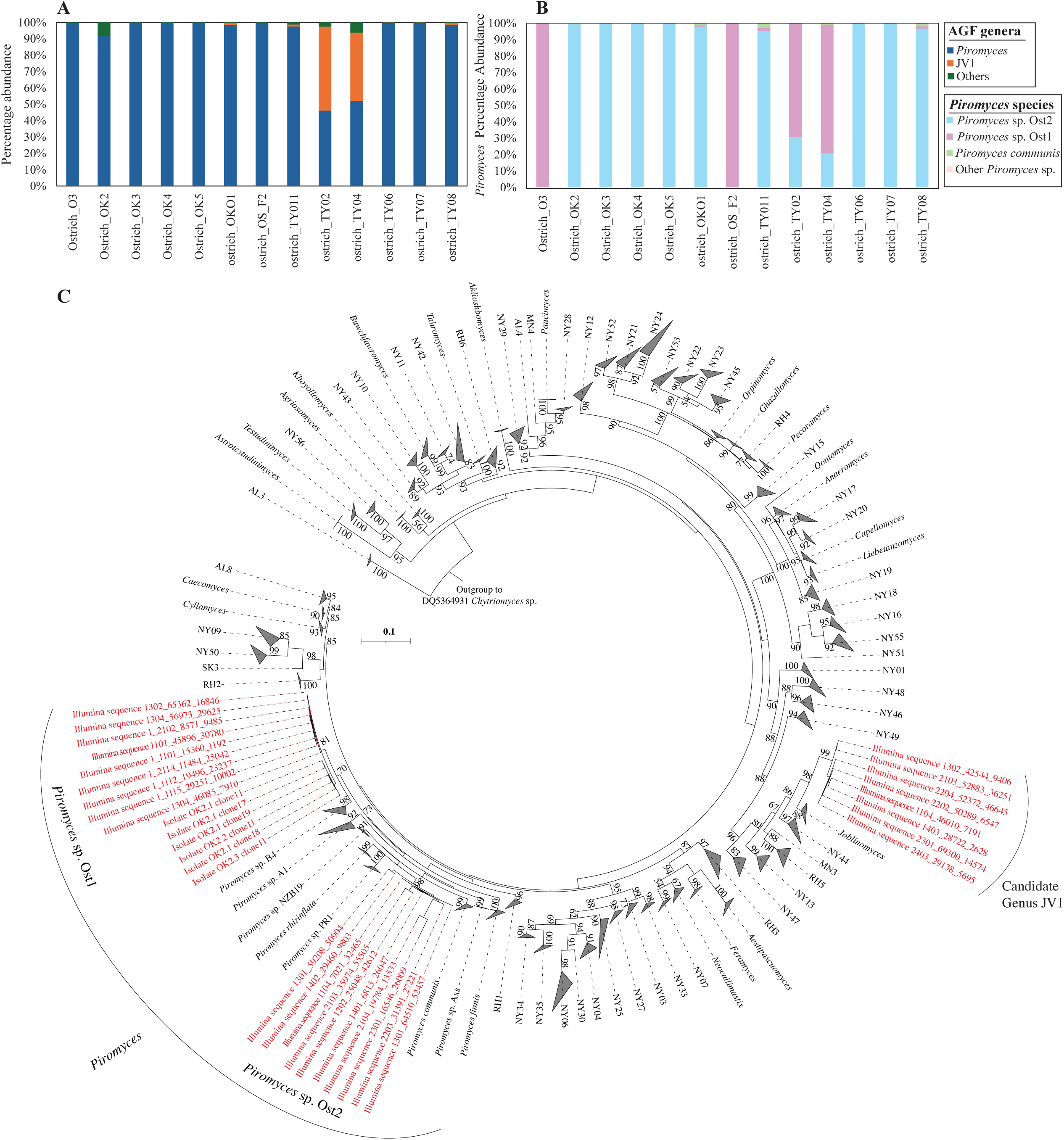
*Neocallimastigomycota* community in ostrich fecal samples. **(A)** Percentage abundance of AGF genera in ostrich fecal samples. “Others” includes all other genera identified outside of *Piromyces* and the putative novel genus JV1 (detailed in Table S1). **(B)** Putative species-level affiliation of sequences belonging to genus *Piromyces* in ostrich fecal samples. **(C)** Phylogenetic tree depicting the position of the two novel *Piromyces* species (*Piromyces* sp. Ost1 and *Piromyces* sp. Ost2) as well as the novel candidate genus JV1 (in red) in relation to other cultured and uncultured AGF genera.

*Piromyces* sp. Ost2 was more frequently encountered in mammalian (404 samples out of 661)^6^, marsupial (49 samples out of 61)^9^, and tortoise (7 samples out of 11)^11^ samples. However, while dominant in ostrich samples, *Piromyces* sp. Ost2 always represented a small fraction of the overall community in other hosts (0.55% of the total community in mammals^6^, 0.239% of the total community in marsupials^9^, and 2.2% of the total community in tortoises^11^) (Table 2).

**Table 2.** Ecological distribution of the ostrich specific *Piromyces* species and uncultured candidate genus JV1.

|  | Closest cultured representative |  | Occurrence in previous studies (sequences with % similarity >97%) |  |  |  |  |  |  |  |
| --- | --- | --- | --- | --- | --- | --- | --- | --- | --- | --- |
|  | Genus | % similarity | Mammals<sup>8</sup><br>(661 samples, 8,772,160 sequences) |  | Mammals<sup>24</sup><br>(12 samples, 304, 958 sequences) |  | Marsupials<sup>11</sup><br>(61 samples, 174,959 sequences) |  | Tortoises<sup>13</sup><br>(11 samples, 40,413 sequences) |  |
|  |  |  | Number | % | Number | % | Number | % | Number | % |
| *Piromyces* sp. Ost1 | *Piromyces* sp A | 96.55 | 0 | 0 | 0 | 0 | 2 | 0.001 | 8 | 0.020 |
| *Piromyces* sp. Ost2 | *P. communis* clone P | 95.25 | 48556 | 0.554 | 0 | 0 | 419 | 0.239 | 876 | 2.168 |
| JV1 | *Joblinomyces apicalis* | 93.63 | 762 | 0.009 | 0 | 0 | 39 | 0.022 | 1 | 0.002 |

In two out of 13 ostrich samples, roughly half the community encountered was neither affiliated with the genus *Piromyces* nor with any of the currently recognized AGF genera^4,5^ and candidate genera^6,57,58^. Rather, it belonged to a monophyletic novel genus-level clade, to which the name JV1 is proposed (Figure 1A). Sequence divergence within the JV1 clade was low (0.28-1.4%), indicating that all JV1 sequences identified constitute a single species. Candidate genus JV1 is most closely related to the genus *Joblinomyces*, exhibiting only 93.63% sequence similarity. Phylogenetic analysis (Figure 1C) confirmed JV1’s position as member of a clade comprising *Joblinomyces* as well as several yet-uncultured AGF genera (NY44, MN3, RH5, NY13, and NY47)^6,57,58^. Assessment of the occurrence of candidate genus JV1 in prior AGF culture-independent diversity surveys with broad host range^6,9,12,22^ indicates that JV1 was occasionally encountered (50/661 of mammalian samples^6^, 4/61 of marsupial samples^9^, and 1/11 of tortoise samples^11^). However, like *Piromyces* sp. Ost1 and Ost2, JV1 sequences always represented a very small fraction of the overall community in these hosts (0.009% in mammalian hosts^6^, 0.022% in marsupial hosts^9^, and 0.002% in tortoise hosts^11^, Table 2).

### Alpha diversity estimates

Ostriches harbored an AGF community with low levels of alpha diversity. On average, 10.31 ± 12.53 genera were encountered per sample, and values of 0.21 ± 0.34 Shannon index, 0.11 ± 0.2 Simpson, and 1.22 ± 0.46 Inverse Simpson were observed (values are average ± SD from the 13 ostrich samples) (Figures 2, S2). These estimates were significantly lower than alpha diversity values in cattle (Wilcoxon P-value < 9.9 x 10^-6^), deer (Wilcoxon P-value < 1.3 x 10^-6^), sheep (Wilcoxon P-value < 1.2 x 10^-5^), goat (Wilcoxon P-value < 9.6 x 10^-7^), and horses (Wilcoxon P-value < 2.8 x 10^-6^), but comparable to values observed in tortoises (Wilcoxon P-value > 0.08), where the AGF community was similarly shown to be dominated by few genera^11^ (Figures 2, S2).

**Figure 2.**
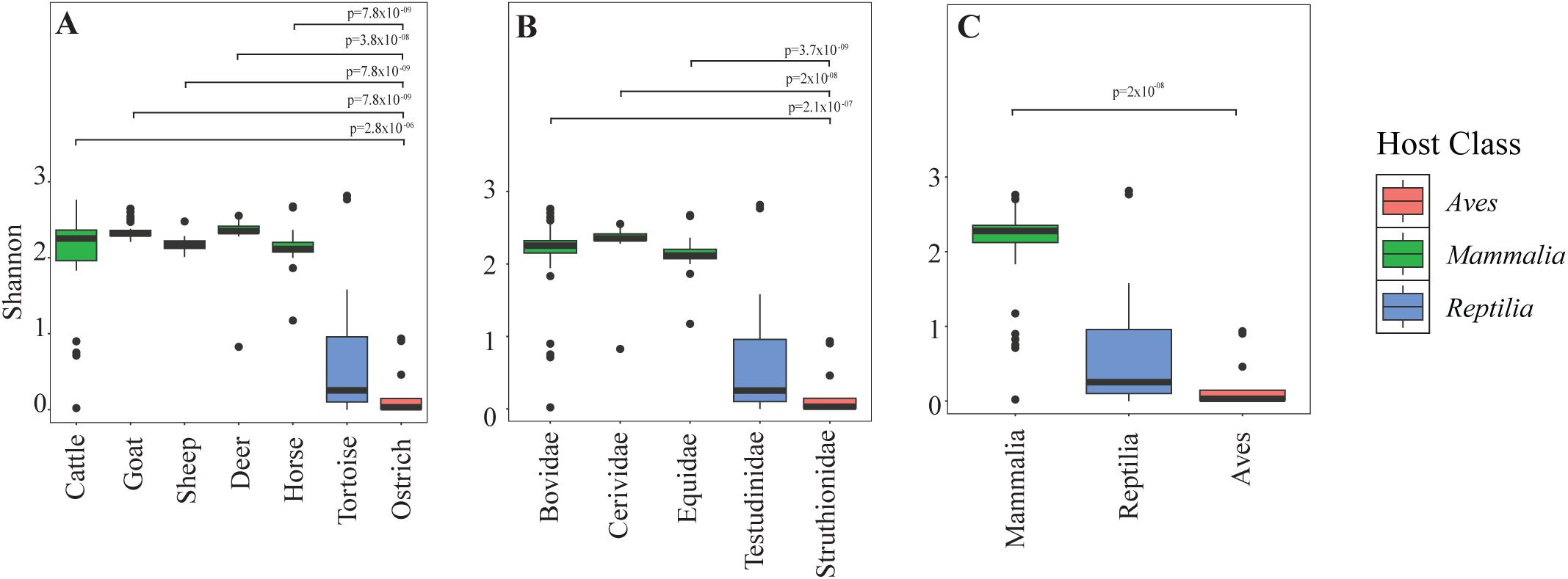
Alpha diversity of *Neocallimastigomycota* in ostriches. Boxplots showing the distribution of Shannon diversity index in ostriches (▪) compared to selected mammalian (▪) and tortoise (▪) samples. Samples were grouped by animal species (A), animal family (B), and animal class (C). Wilcoxon test p-values indicate the significance of differences between ostriches and other mammals. No significant difference (p >0.05) was identified between ostrich and reptilian samples.

### Community structure

AGF community structure in ostriches was compared to that observed in well-studied (cattle, deer, sheep, goat, horses), as well as recently discovered (tortoises) AGF hosts using PCoA constructed based on the phylogenetic similarity-based beta diversity index weighted Unifrac (Figure 3). The first two axes explained 84.3% of the variance. Analysis showed that the animal host species (Figure 3A), family (Figure 3B), and class (Figure 3C) significantly explained 66.57%, 60.81%, and 59.09% of the variance, while the gut type (Figure 3D) only explained 11.05% of the variance. To identify the specific association between AGF genera and ostriches, double PCoA plots were constructed using Bray-Curtis beta diversity indices (Figure 3E) and showed the genus *Piromyces* to be associated with ostriches. Similarly, both LEfSe and Metastats analyses showed the genus *Piromyces* to be differentially abundant in ostriches (LEfSe LDA score of 5.6 and p-value = 0, Metastats p-value = 0.001), and all global phylogenetic signal statistics identified significant correlation between *Piromyces* and ostriches as a host (p-value = 0.001) (Table S2). Finally, LIPA analysis confirmed the strong significant association between *Piromyces* and ostriches (Table S2) (average LIPA value of 7.82, p-value = 0.001). In addition to *Piromyces*, the new uncultured genus JV1 was also differentially abundant in and strongly associated with ostriches (LEfSe LDA score of 4.53 and p-value =4.6 x10^-9^, Metastats p-value =0.001), with significant global phylogenetic signal statistics (p-value <0.002), and high LIPA values in the two ostrich samples in which it was detected (average LIPA=5.52, p-value=0.001) (Table 2).

**Figure 3.**
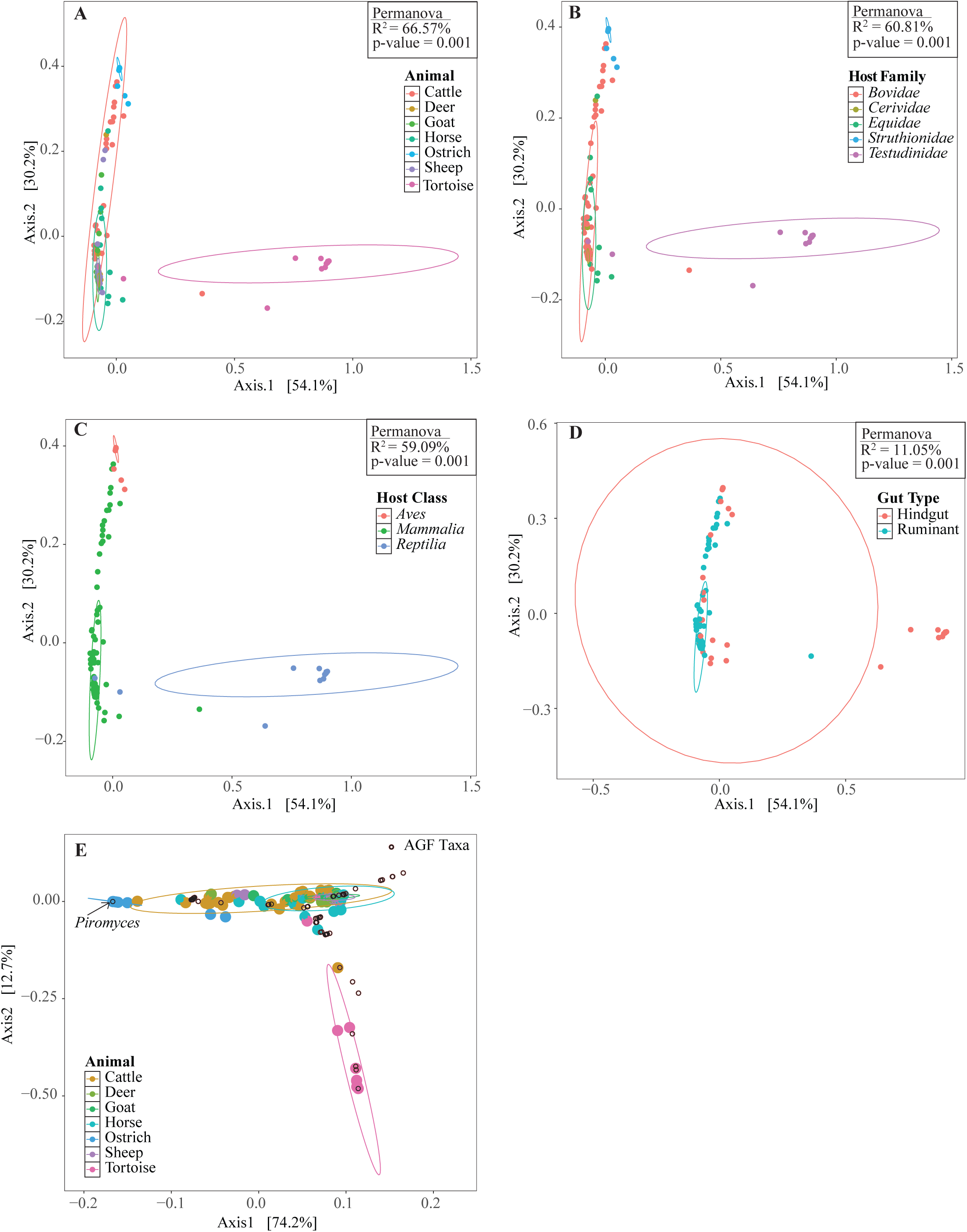
Community structure of *Neocallimastigomycota* in ostriches. (A-D) PCoA plots constructed using weighted UniFrac beta diversity estimates, with a color scheme based on host animal species (A), host family (B), host class (C), and host gut type (D). The % variance explained by the first two axes is displayed on the axes, and results of PERMANOVA for the contribution of host factors to the community structure are shown for each plot (R^2^: the % variance explained by each factor, p: F-test p-value). (E) Double principal coordinate analysis plot constructed using Bray-Curtis beta diversity indices. The AGF taxa are shown as open black circles, and the genus *Piromyces* position is shown with an arrow. Samples are color-coded by the host animal species as in (A).

### Enrichments and isolation of AGF from ostriches

Multiple enrichments were set up, using different media, carbon sources, as well as different incubation temperatures (Table 1). Successful enrichment efforts yielded visual biomass, gas bubbles, and clumping and floating of plant biomass (when used), with the identity of AGF determined to be either *Piromyces* sp. Ost1 or candidate genus JV1 by PCR amplification (D1-D2 region of the LSU) and Sanger sequencing. Repeated isolation efforts yielded multiple representatives of *Piromyces* sp. Ost1 (7 strains). Transcriptomic sequencing and subsequent phylogenomic analysis (Figure 4) confirmed the position of *Piromyces* sp. Ost1 as a novel species within the genus *Piromyces*.

**Figure 4.**
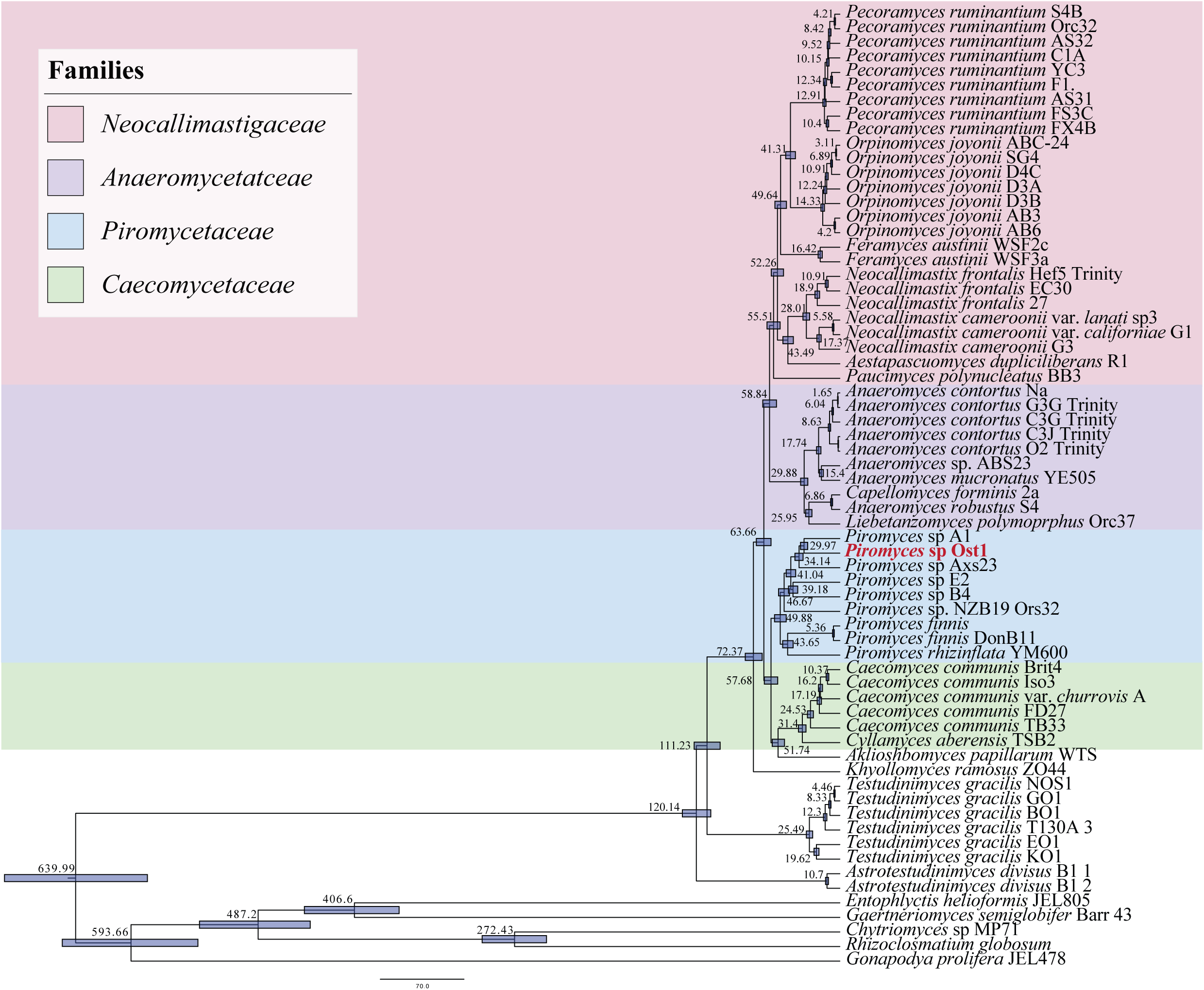
Phylogenomic analysis and molecular timing of strain Ost1. Bayesian phylogenomic maximum clade credibility (MCC) tree of *Neocallimastigomycota* with estimated divergence time for major nodes. Estimate for the divergence time of *Piromyces* sp. Ost1 from its closest mammalian relative (*Piromyces* species A1) is highlighted. The 95% highest probability density (HPD) ranges (blue bars) are denoted on the nodes, and the average divergence times are shown.

Despite repeated attempts, no pure culture of the candidate genus JV1 from positive enrichments could be obtained. Furthermore, *Piromyces* sp. Ost2 was never enriched (Table 1), despite its predominance in many samples (Figure 1B).

### Timing the evolution of *Piromyces* sp. Ost1

Transcriptomics-enabled molecular clock timing suggested a divergence time estimate of ≈ 30 Mya (95% highest probability density interval of 26.95-32.94 Mya) for *Piromyces* sp. Ost1 (Figure 4). Such time postdates the evolution of the infraclass *Palaeognathae* (∼72.8-110 Mya)^59,60^, comprising the flightless birds and the volant tinamous, as well as the diversification of *Struthioniformes* and the genus *Struthio* (∼69-79.6 Mya)^60,61^, but might have coincided with the evolution of flightlessness in these lineages.^59^

### Comparative gene content and CAZyome analysis

Comparative genomic analysis demonstrated broadly similar COG, KOG, and KEGG profiles between *Piromyces* sp. Ost1 and AGF obtained from mammalian hosts (Figure 5). PCoA based on GH family composition for *Piromyces* sp. Ost1 grouped *Piromyces* sp. Ost1 (Figure 5, grey triangle) with the mammalian AGF (Figure 5, circles, n = 53). Both groups were distinct from tortoise AGF (n = 7), which were previously shown to possess a unique and highly reduced CAZyme repertoire.^11^ Overall, comparative gene content and CAZyome analysis suggest functional similarity between ostrich-sourced and mammalian-sourced AGF.

**Figure 5.**
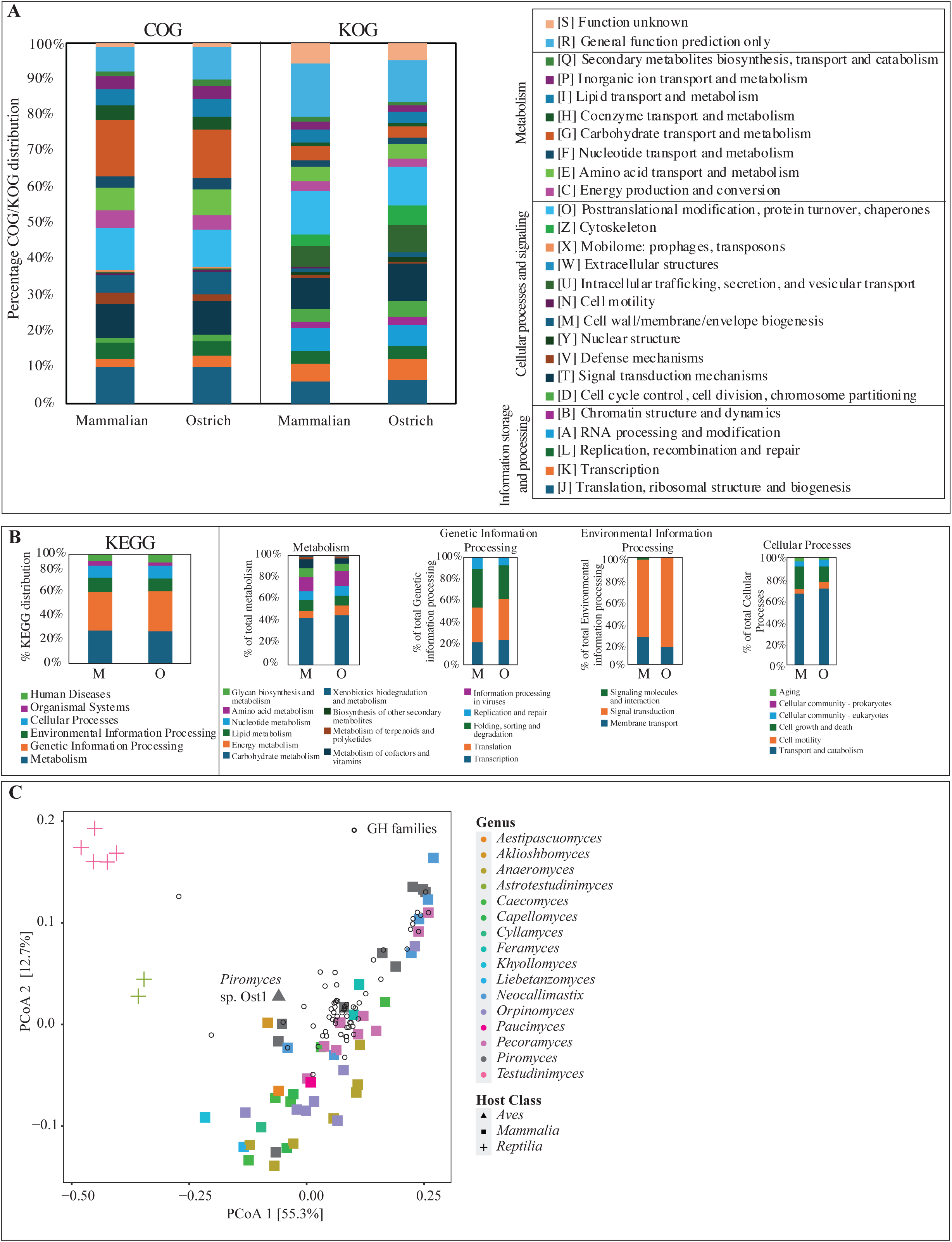
Comparative gene content analysis. (A-B) Gene content comparison between mammalian sourced (M; left stacked columns) and *Piromyces* sp. Ost1 sourced (O; right stacked columns) transcriptomes using COG/KOG (A), and KEGG (B) classification. KEGG classification is further broken down into four main categories: Metabolism, Genetic Information Processing, Environmental Information Processing, and Cellular Processes. (C) Principal coordinate analysis (PCoA) biplot based on the GH families’ composition in *Piromyces* sp. Ost1 transcriptome (grey triangle) compared to 67 previously obtained AGF transcriptomes belonging to 16 genera (including the genus *Piromyces*). The % variance explained by the first two axes is displayed, and strains are color-coded by AGF genus, as shown in the figure legend. The shapes correspond to the host class, with mammals shown as ▪, *Aves* shown as ▲, and reptiles shown as “+”. GH families are shown as empty circles with black borders.

## Discussion

Our investigation of AGF in ostriches revealed a uniform (Figure 1A), low diversity community (Figure 2, Table S1) that was mostly comprised of novel AGF taxa (Figure 1A, 1B). The ostrich AGF community was distinct from previously described AGF communities (Figure 3), with ostrich-associated taxa rarely encountered in mammalian, marsupial, or tortoise datasets (Table 2).

Sequences putatively representing two novel species within the genus *Piromyces* constituted the majority of the AGF community in eleven out of thirteen ostrich samples and roughly half the community in the remaining two (Figure 1A, 1B). The genus *Piromyces* is ubiquitous, representing an integral member of the AGF diversity in a wide range of mammalian foregut and hindgut fermenters. *Piromyces* was one of the earliest AGF genera to be identified^62^, isolated^63^, named^55^, and characterized.^62^ Historically, thallus morphology and flagellation of zoospores were used for taxonomic characterization of AGF isolates, and the clade *Piromyces* comprised any strain with filamentous rhizoids, monocentric thallus development, and monoflagellated zoospores. Currently, the genus *Piromyces* includes all isolates phylogenetically affiliated with the first described monocentric, monoflagellated and filamentous isolate (*Piromyces communis*)^4,55,62–64^. A recent large-scale analysis of available sequencing data for *Neocallimastigomycota* concluded that current members of the genus *Piromyces* display a higher level of within-genus sequence divergence in marker genes (e.g., D1-D2 LSU ranging from 1.24-5.6% with an average of 3.4%), as well as in whole genome metrics (AAI ranging from 72.58–99.06% with an average of 79.35%) than that typically encountered within other genera (genus cut-off set at 3% sequence divergence in D1-D2 LSU and 85% AAI).^4,35^ While these values support its breakdown into multiple genera, the genus was retained as a single entity^4^, partly due to the lack of sequence data from now extinct original type strains for *Piromyces* species (e.g., *P. mae*, *P. dumbonicus*, *P. minutus*, *P. spiralis*, and *P. citronii*)^4,65,66^

The two novel, ostrich-associated species encountered in this study exhibited large sequence divergence values from their closest relatives (3.45% and 4.75% D1-D2 LSU sequence divergence for sp. Ost1 and sp. Ost2, respectively). These values would have justified their placement as a new genus had they belonged to a different clade/family within the *Neocallimastigomycota*. While it is possible that both novel species identified in this study could belong to previously described *Piromyces* species lacking sequence data, this seems unlikely given that all described species of *Piromyces* have been isolated from mammalian hosts^4^, whereas these ostrich-associated species have rarely been identified in mammals (Table 2).

Two out of 13 ostrich samples were dominated by a genus-level cluster (JVI) whose closest relatives (93.63% LSU sequence similarity) belong to the genus *Joblinomyces* (Table 2, Figure 1C, Table S1). Both samples came from the same ostrich farm in Texas, U.S.A. (Table S1). All samples from this farm contained sequences affiliated with JV1 (0.18 – 49.65%), while only one sample (out of seven) from other farms/zoos harbored JV1. Given the relatively low number of samples (n = 13) and locations (n = 4) investigated in this study, an accurate assessment of the global prevalence pattern of candidate genus JV1 in ostriches is not feasible. Still, examination of the ecological distribution of candidate genus JV1 in previously published mammalian, marsupial, and reptilian datasets revealed its extremely low abundance (Table 2), hinting that this strain could either be very rare in general or ostrich specific. While JV1 could not be isolated in pure culture in this study, it was enriched at two different temperatures (39 ℃ and 41 ℃), and repeated efforts for its isolation are ongoing (Table 1).

The AGF community in ostriches exhibited low levels of alpha diversity, driven by the predominance of one genus (i.e., >50% relative abundance) in most samples. In a meta-analysis of data published in three studies^6,9,13^, such a pattern of predominance was also observed in 82% of investigated tortoises (9/11 samples)^13^ and in 37% of investigated mammals (264 out of 722 samples, Figure 6B, Table S3).^6,9^ Ostriches and tortoises are hindgut fermenters, and a pattern of predominance has previously been linked to hindgut fermenters in mammals^6^. This link between predominance and hindgut fermentation is corroborated by our study (100% of ostrich samples were dominated by one genus) as well as the meta-analysis of previously published results, where 25% of foregut and 64% hindgut fermenter samples showed this pattern (Table S3). Furthermore, it is noteworthy that the data is heavily biased, with 74% of total samples obtained from cattle, horses, donkeys, goats, and sheep. Of the hindgut fermenter samples, 63% were obtained from horses and donkeys, while the foregut fermenter samples were mainly sourced from cattle, goats, and sheep (79% of total foregut fermenter samples).

**Figure 6.**
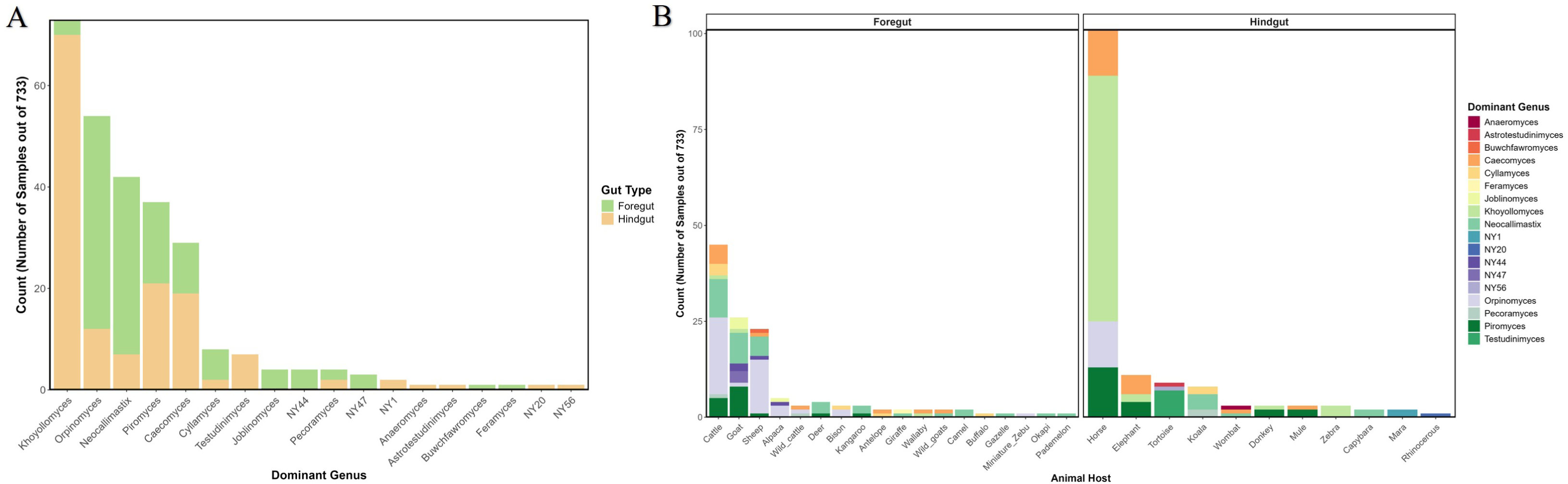
A meta-analysis assessing AGF predominance patterns. Data from three previously published amplicon sequencing studies (targeting the same marker gene region and using the same bioinformatic analysis pipeline)^6,9,13^ were used to assess patterns of AGF predominance (i.e., a single genus representing >50% of the AGF community) within various hosts. (A) Predominant genera and the gut type they are associated with. (B) The absolute number of samples per animal host that showed a predominance pattern and the AGF genera associated with it. For details on the meta-analysis, refer to Table S3.

While the genus *Piromyces* predominated in ostrich samples in this study, other genera were observed to dominate samples in prior studies. In three studies (a total of 733 samples), the following genera predominated: *Khoyollomyces* (n = 73), *Orpinomyces* (n = 54), *Neocallimastix* (n = 42), *Piromyce*s (n = 37), and *Caecomyces* (n = 29) (Figure 6A). *Khoyollomyces* and *Caecomyces* primarily dominate mammalian hindgut fermenters (e.g., horses, elephants), while *Orpinomyces* and *Neocallimastix* are more common in mammalian foregut fermenters (e.g., cattle, sheep, goats) (Figure 6B, Table S3). *Piromyces* showed no clear preference, dominating foregut (n = 16) and hindgut (n = 21) fermenters. The reasons underpinning the ecological success of some AGF genera over others are currently unclear. More studies to identify metabolic, physiological, and genomic differences between various AGF taxa, as well as linking such differences to observed host and gut-type preferences of AGF genera are sorely needed to address such issues.

Our results show a clear pattern of host-AGF preference in ostriches (Figure 3, Table S2). Our molecular timing analysis estimated an evolutionary time for *Piromyces* sp. Ost1 of ≈ 30 Mya. In the context of bird evolution, it is known that birds first appeared in the fossil record during the Middle-Late Jurassic (∼165-150 Mya), diversified by the early Cretaceous, with true modern birds radiating post-Cretaceous and surviving the Cretaceous-Paleogene (K-Pg) extinction event.^67^ Within extant birds, the *Palaeognathae* (which includes the flightless ratites and the tinamous) diversified first (72.8-110 Mya)^59–61^, followed by the diversification of *Struthioniformes* (∼69-79.6 Mya).^60,61^ Flightlessness evolved in *Struthioniformes* around 25-30 Mya, and the process was tightly associated with the development of herbivory.^59^ We therefore propose that the evolutionary timeline of *Piromyces* sp. Ost1 aligns with the emergence of flightlessness and herbivory within the ancestors of modern ostriches. This suggests a pattern of co-evolution and subsequent retention throughout time up to the evolution of modern ostriches (estimated evolution at 5.3-2.6 Mya).^68^ An alternative scenario, where ostrich specific *Piromyces* sp. Ost1 evolved independently in an unknown host before colonizing modern ostriches after their speciation, cannot be ruled out. However, we deem this scenario less plausible since *Piromyces* sp. Ost1 was rarely identified in animals outside the *Palaeognathae* (mammals or reptiles) and appears to be specific to the ostrich alimentary tract (Table 2), potential sampling biases notwithstanding.

Finally, to investigate why *Piromyces* sp. Ost1 is highly successful in ostriches but unable to effectively colonize other AGF hosts, we conducted a transcriptomic analysis comparing AGF sourced from different hosts. Comparative gene content analysis showed similar functional profiles (COG, KOG, and KEGG) in *Piromyces* sp. Ost1 compared to mammalian-sourced AGF taxa (Figure 5A and 5B). *Piromyces* sp. Ost1 showed a similar CAZyome to mammalian AGF taxa, including mammalian *Piromyces* species (Figure 5C). While more detailed analysis could clarify finer levels of substrate utilization patterns, the lack of stark differences in broad plant biomass degradation capacities compared to mammalian-sourced isolates is noted. We therefore hypothesize that physiological differences in the ostrich gut (e.g., a slightly higher temperature of 38.1°C to 40.5°C) ^69,70^ compared to mammalian hindgut fermenters (e.g., 37.5 °C to 38.5 °C in horses)^71^ could play an important role in the selection of this specific *Piromyces* species. Our preliminary analysis of *Piromyces* sp. Ost1 showed tolerance to higher temperatures and a broader temperature growth range compared to mammalian-sourced *Piromyces* (unpublished data). It is also possible that AGF in the ostrich gut have additional roles beyond plant biomass breakdown, e.g., detoxification, or secondary metabolites secretion, with ostrich AGF specifically adapted to play this role.^72,73^ A more detailed, in-depth experimental and omics-based investigation is needed to address such an interesting question.

In conclusion, this study has expanded the known host range for AGF to include birds (class *Aves*), specifically the common ostrich (*Struthio camelus*). We demonstrate a strong pattern of AGF-host specificity in ostriches and suggest that such a pattern has arisen out of co-evolutionary phylosymbiosis. However, it is important to note that the ostriches investigated in this study were limited to domesticated animals within the south-central part of the United States. Future studies on wild ostriches, as well as other hindgut fermenting animals outside *Mammalia* are needed to confirm the results observed and expand on the global diversity and the evolutionary patterns in *Neocallimastigomycota*.

## Acknowledgments

Work in M. S. Elshahed and N. H. Youssef Laboratories was supported by the United States National Science Foundation (NSF) grant number 2029478, and the United States National Institute of Health (NIH) grant number P20GM152333-01. We thank the Oklahoma City Zoo, Happy Acres Ostrich Ranch LLC, Snider Family Exotics, and Haley Anthony for providing fecal samples. We would further like to thank Kale Goodwin for his efforts in isolating AGF from ostrich fecal samples and the Boren Veterinary Medical Teaching Hospital for providing rumen fluid.

## Author contributions

Fund acquisition, project supervision: MSE and NHY. Sample collection: MSE, ALJ, CHM, CJP. Lab work: JV, KN, ALJ, TY, CHM, CJP. Data analysis: NHY, JV. Manuscript: MSE, NHY, JV. Figures: NHY and JV.

