## Supplementary Figure 1, Table S2 for "The anaerobic gut fungal community in ostriches (*Struthio camelus*)"

Supplementary Tables (provided as an Excel sheet)

**Table S1.** A list of the 13 samples studied, their sources, the total number of sequences analyzed per sample, sample coverage, and the percentage abundance of AGF genera in each sample.

**Supplementary Table S3.** Comparative analysis of *Neocallimastigomycota* community structure (genus level) in various hosts. The meta analysis is based on the OTU count tables from amplicon sequencing with *Neocallimastigomycota* specific primers targeting the LSU region taken from previously published studies (Meili *et al.* 2023, Jones *et al.* 2024, Pratt *et al.* 2024).

**Table S2.** Results of Metastats, linear discriminant analysis (LDA) effect size (LEfSe), global, and local phylogenetic signal statistics for the significant association of *Piromyces* and candidate genus JV1 with ostriches.

| Genus | Metastats | | | | LEfSe | | Global phylogenetic signal | | | | | | Local phylogenetic signal (LIPA) | |
|---|---|---|---|---|---|---|---|---|---|---|---|---|---|---|
| | Average ±SD in Aves | Average ±SD in Mammalia | Class of significant enrichment | p-value | LDA value | p-value | Abouheif Cmean | | Moran I | | Pagel Lambda | | | |
| | | | | | | | | p | | p | | p | LIPA | p |
| *Piromyces* | 90.93±19.2 | 10.96±12.2 | Aves | $9.9x10^{-4}$ | 5.603 | 0 | 0.72 | 0.001 | 0.765 | 0.001 | 0.812 | 0.001 | 7.8±1.96 | 0.001 |
| JV1 | 7.3±16.99 | 0 | Aves | $9.9x10^{-4}$ | 4.53 | $4.67x10^{-9}$ | 0.078 | 0.002 | 0.083 | 0.001 | 0.19 | 0.001 | 5.52±0.055* | 0.001 |

*Values are averages ± Standard deviation in the two ostrich samples with >40% abundance of JV1. When all samples were considered, the average LIPA was 0.79±2.1.

Supplementary figures.

**Figure S1. Alpha diversity of *Neocallimastigomycota* in Ostriches**. Boxplots showing the distribution of Observed number of genera (A, D, G), Simpson diversity index (B, E, H), and Inverse Simpson (C, F, I) in ostriches (■) compared to selected mammalian (■) and tortoise (■) samples. Samples were grouped by animal species (A-C), animal family (D-F), and animal class (G-I). Wilcoxon test p-values are shown for the significance of difference between ostriches and other mammals. No significant difference ($p > 0.05$) was identified between ostrich and reptilian samples.

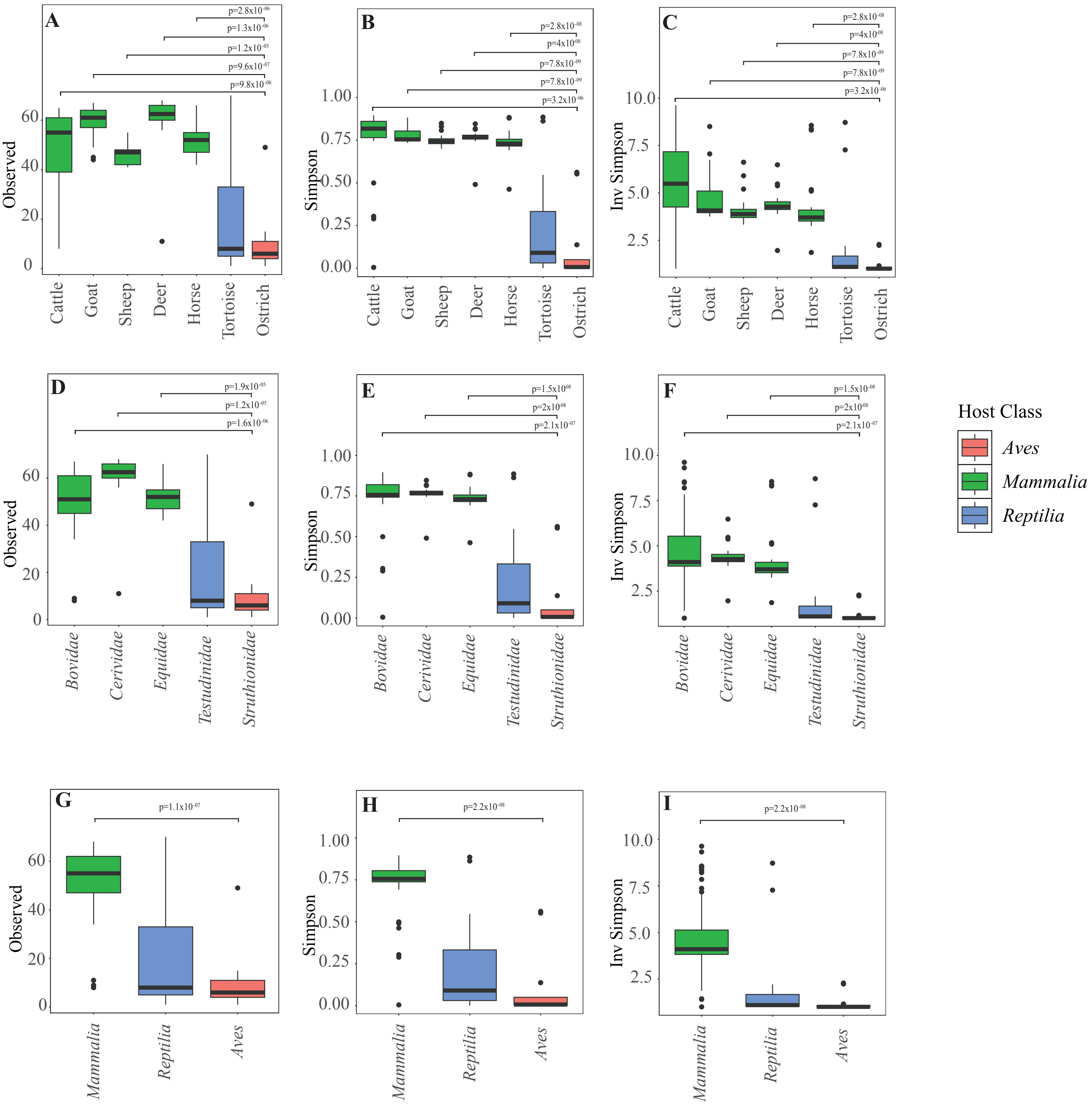

A
B
C
D
E
F
G
H
I
Observed
Simpson
Inv Simpson
p=2.8x10-06
p=1.3x10-06
p=1.2x10-05
p=9.6x10-07
p=9.8x10-06
p=2.8x10-08
p=4x10-08
p=7.8x10-09
p=7.8x10-09
p=3.2x10-06
p=1.9x10-05
p=1.2x10-05
p=1.6x10-06
p=1.5x10-08
p=2x10-08
p=2.1x10-07
p=1.1x10-07
p=2.2x10-08
Cattle
Goat
Sheep
Deer
Horse
Tortoise
Ostrich
Bovidae
Cerividae
Equidae
Testudinidae
Struthionidae
Mammalia
Reptilia
Aves
Host Class
Aves
Mammalia
Reptilia
